# The structural basis of itch receptor MRGPRX1 activation

**DOI:** 10.1101/2022.12.21.521535

**Authors:** Bing Gan, Leiye Yu, Haifeng Yang, Haizhan Jiao, Bin Pang, Yian Chen, Chen Wang, Rui Lv, Hongli Hu, Zhijian Cao, Ruobing Ren

## Abstract

Mas-related G-protein-coupled receptors X1-X4 (MRGPRX1-X4) are four primate-specific receptors that are recently reported to be responsible for many biological processes, including itch sensation, pain transmission, and inflammatory reactions. MRGPRX1 is the first identified human MRGPR, and its expression is restricted to primary sensory neurons. Due to its dual roles in itch and pain signaling pathways, MRGPRX1 has been regarded as a promising target for itch remission and pain inhibition. Here, we reported a cryo-electron microscopy (cryo-EM) structure of G_q_-coupled MRGPRX1 in complex with a synthetic agonist compound 16 in an active conformation at an overall resolution of 3.0 angstrom via a NanoBiT tethering strategy. Compound 16 is a new pain-relieving compound with high potency and selectivity to MRGPRX1 over opioid receptor. According to the structure analysis, we revealed that MRGPRX1 shares common features of the G_q_-mediated receptor activation mechanism of MRGPRX family members. However, the variable residues in orthosteric pocket of MRGPRX1 exhibits the unique agonist recognition pattern, which may facilitate to design MRGPRX1-specific modulators. Together with receptor activation and itch behavior evaluation assays, our study provides a structural snapshot to modify therapeutic molecules for itch relieving and analgesia targeting MRGPRX1.

## Introduction

Itch is defined as the sensation that causes the desire to scratch the skin[1]. It is a common and frequently occurring symptom associated with many skin diseases among humans[2]. Numerous factors can induce itches, such as chemicals, insect bites, and even self-generated substances resulting from varies diseases[3]. Unfortunately, due to diverse inducements and complicated pathogenesis, treating itch in the clinic is still challenging, especially the chronic itch, which will devastate people and cause much suffering[4]. The itch can be generally divided into histaminergic and nonhistaminergic[5]. Usually, most histaminergic itch results in acute itch, whereas chronic itch is more probable to be nonhistaminergic[6]. Therefore, the well-developed antihistamine drugs are inefficient in chronic itch relieving, which suggests the significance of finding novel drug targets for chronic itch treatment[6].

Mas-related G-protein-coupled receptors (MRGPRs) have been recently identified as pruritogenic receptors mediating the non-histaminergic itch[7]. The *Mrgpr* gene family encodes MRGPRs, a large family which comprises 27 and 8 members in mice and humans, respectively[7, 8]. MRGPRX1-X4 are four primate-specific receptors, suggesting that the X subfamily may be a simplified alteration in human evolution[4]. MRGPRX1 is the first identified human MRGPR that expresses in dorsal root ganglia (DRG) and trigeminal ganglia (TG) specifically[4]. Compared with other MRGPRX members, MRGPRX1 stands out for its dual roles in mediating itch[9] and inhibiting persistent pain[10]. Persistent pain is a severe health problem worldwide, and ordinary analgesics like opioids targeting opioid receptors may lead to several side effects such as drug addiction[10, 11]. Notably, MRGPRX1 is insensitive to the classical opioid receptor antagonists, indicating that MRGPRX1 could be a new target for treating chronic pain[10].

Extensive studies of MRGPRX1 were conducted in inflammation, pain and itch sensations[7]. A series of natural and synthetic agonists, antagonists, and allosteric modulators of MRGPRX1 have been developed[12]. However, there is currently no drug targeting MRGPRX1 commercialized. The structure determination of GPCR may provide the detailed molecular basis of ligand interaction to facilitate modulator development[13]. Recently, two groups reported the agonist-stabilized cryo-EM structures of MRGPRX2[14, 15] and MRGPRX4[14] in complexes with trimeric G proteins. The structural characteristics of orthosteric pockets and modulator specificities are examined thoroughly. The critical acidic residues D184^5.38^ and E164^4.60^ in MRGPRX2 and the entirely positive orthosteric pocket in MRGPRX4 mainly determines the chemical property of varies modulators.

In this study, we reported the cryo-EM structure of the active MRGPRX1-G_q_ complex bound to compound 16 at an overall resolution of 3.0 angstrom. Compound 16 is a new synthetic MRGPRX1 agonist with high potency and selectivity over other MRGPRXs and opioid receptor[16]. Our complex structure reveals the conserved mechanism of small molecule-induced receptor activation among the MRGPRX receptors. The structure also clearly presents a highly conserved orthosteric pocket for natural agonist recognition, such as bovine adrenal medulla 8-22 peptide (BAM8-22)[17], γ2-MSH[18, 19], and conopeptide (CNF-Tx2)[20]. Notablly, a few variable residues in orthosteric pocket of MRGPRX1 exhibit the unique agonist recognition pattern for compound 16. These findings will give us clues to the modification of small molecule scaffolds targeting MRGPRX1 specifically and accelerate the development of the novel drug for itch and pain relieving.

## Results

### The overall structure of G_q_-coupled MRGPRX1 bound to compound 16

To improve receptor expression, we fused thermostabilized apocytochrome b562 (BRIL) at the N-terminus of MRGPRX1[21]. NanoBit tethering strategy was used for the complex formation, with the LgBit and HiBit fused to the C-terminus of the receptor and Gβ subunit, respectively[22] (Supplementary Figures 1 A and B). We used Bioluminescence Resonance Energy Transfer (BRET) assay to evaluate the impact of receptor modification on G protein coupling capability. The fusion of BRIL and LgBit to receptor only marginally affected receptor activity (Supplementary Figure 1C, Supplementary Table 1). To further stabilize the complex, we used an engineered G_αq_ chimera in the complex assembly. The G_αq_ chimera was designed based on the mini-G_αs/q71_ scaffold[23, 24]. The N-terminus of G_αq_ was replaced by the N-terminus of G_αi1_. Similar methods had been successfully used in the structure determination of several G_q_-bound GPCRs, including the G_q_-bound 5-HT2A receptor[25] and ghrelin receptor[24]. The engineered G_αq_ chimera used in the further structure study will be simplified as G_αq_. We co-expressed BRIL-MRGPRX1-Lgbit, G_αq_, and G_βγ_-HiBit to obtain the MRGPRX1-G_q_ complex. The complex was further stabilized by incubating with Nb35[26] and scFv16[27] in the presence of compound 16. (Supplementary Figure 2).

The compound 16-MRGPRX1-G_αq_ complex structure was determined by cryo-EM to yield a final map at an overall resolution of 3.0 Å (Figure 1A, Supplementary Figures 3A–F, Supplementary Table 2). In the map, the densities for the receptor, G_αq_, G_βγ_, Nb35, scFv16, and compound 16 could be well distinguished, and the interface residues between MRGPRX1 and G_αq_ (α5-helix) were clearly defined (Supplementary Figure 3G). Thus, we built a reliable atomic model based on the well-traced α-helices and aromatic side chains (Figure 1B). Due to the flexibility, the N-terminus (M1-K25), part of the extracellular loops (I90) and long C-terminal residues (R279-Q322) of the receptor are invisible.

**Figure 1.**
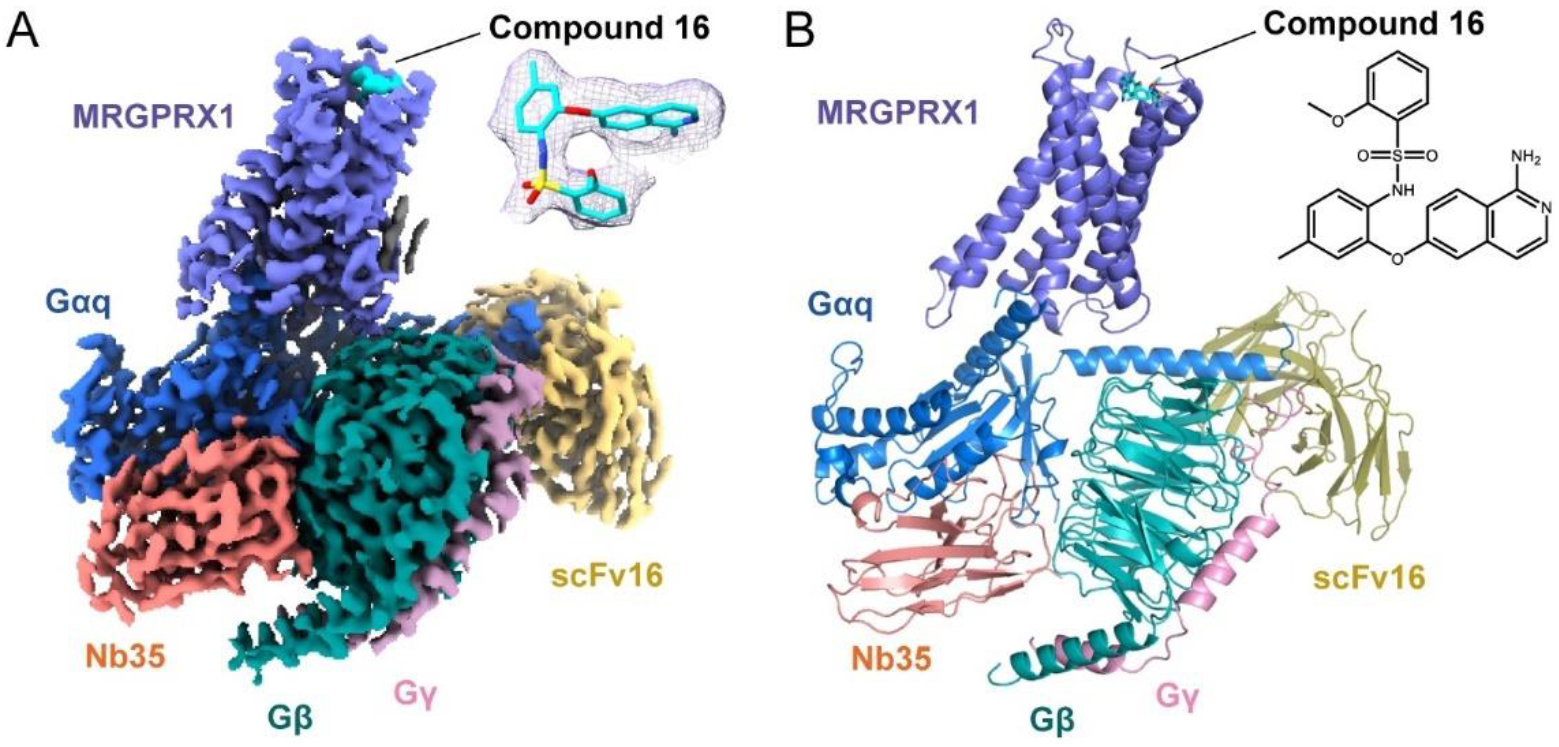
Cryo-EM structure of compound 16-MRGPRX1-G_q_ complex. (A) Cryo-EM density map of MRGPRX1-G_q_ in complex with compound 16. Receptor, compound 16, G_q_, G_β_, G_γ_, Nb35, and scFv16 are colored slate, cyan, marine, forest, pink, salmon, and yellow-orange, respectively.. The density of compound 16 is shown in mesh at the top-right corner. Compound 16 is fitted and shown as sticks. (B) Structural model for the compound 16-MRGPRX1-G_q_ complex. Receptor, ligand, G_q_, G_β_, G_γ_, Nb35, and scFv16 are colored the same as (A). Compound 16 is shown as sticks. The 2D chemical structure of compound 16 is shown in the top-right corner.

### The orthosteric pocket of MRGPRX1

The MRGPRX1 exhibits a shallow, broad, and wide-open ligand binding pocket (Figure 2A). The distance between compound 16 and the critical toggle switch residue G229^6.48^ is about 16.8 Å (Figure 2B). It indicates that compound 16 is positioned near the extracellular surface but not buried deep in the receptor. The shallow pockets are also observed in the MRGPRX2-G_αq_[14, 15] and MRGPRX4-G_αq_ complex[14], suggesting the common pocket features among all MRGPRX receptors. Compound 16 occupies only about one-third of the pocket (Figure 2A) but is sufficient for the receptor activation. Similarly, the agonist (R)-ZINC-3573 and MS47134 take up only a small part of the pocket in MRGPRX2 and MRGPRX4, respectively (Supplementary Figures 4A and B). The structure alignment shows that three agonists bind to different regions of the orthosteric pocket in three receptors, indicating distinct recognition mechanisms (Supplementary Figure 4C). The electrostatic potential of the MRGPRX1 pocket is partially negative, partially positive, and partially hydrophobic (Supplementary Figure 4D). In contrast, the electrostatic potential of the MRGPRX2 pocket is partially negative (sub-pocket 1) and partially hydrophobic (sub-pocket 2), and the electrostatic potential of MRGPRX4 pocket is positive (Supplementary Figures 4E and F). It suggests that MRGPRXs may prefer agonist scaffolds with distinct electro-properties.

**Figure 2.**
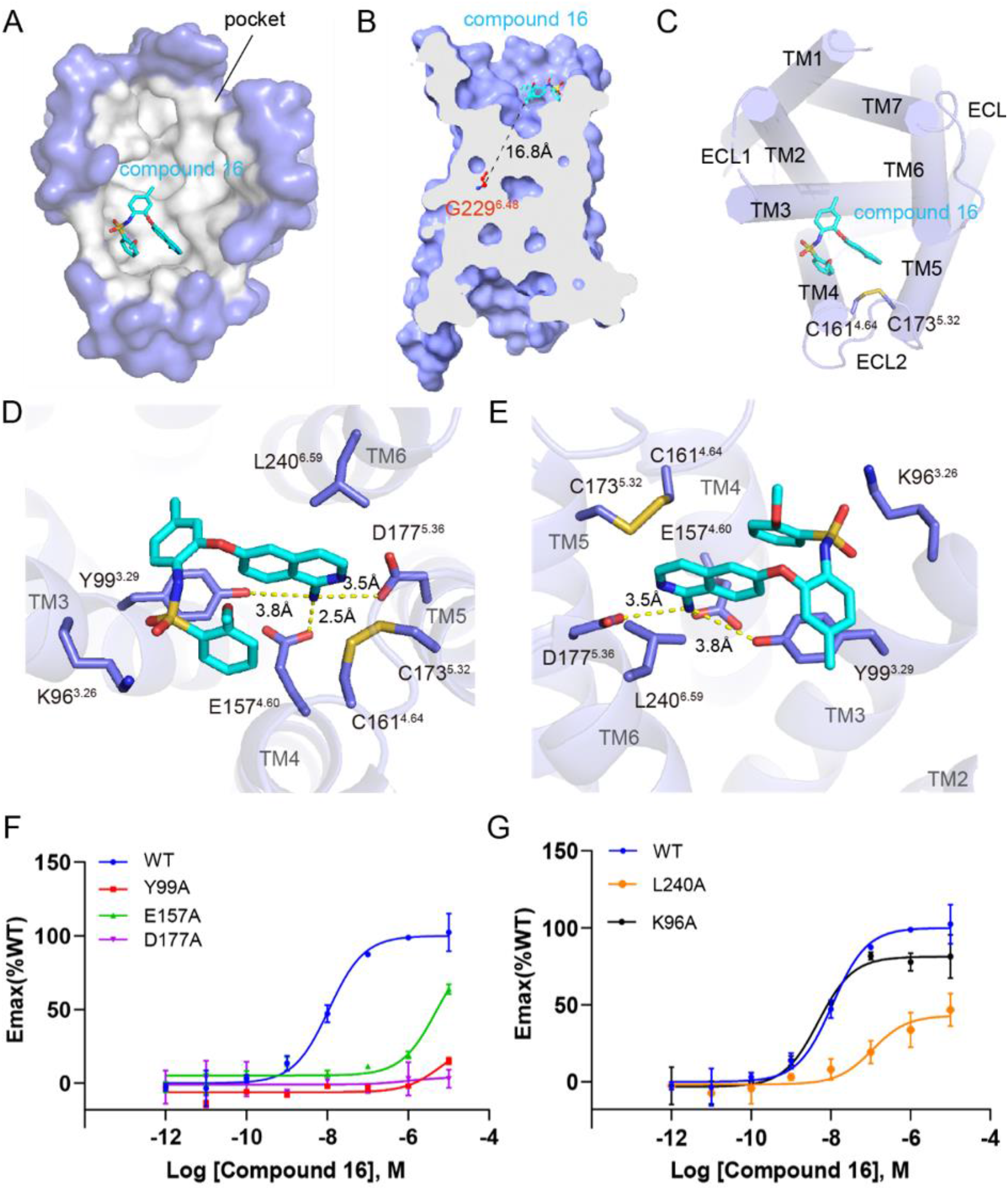
The compound 16 binding pocket in MRGPRX1. (A) Top view of the compound 16 binding pocket from the extracellular side (surface mode). Pocket is colored gray, and compound 16 is shown as cyan sticks. (B) Cut-away view of the compound 16-binding pocket. The distance between the ligand and the traditional toggle switch position is shown as dashed lines. (C) Top view of the compound 16 binding pocket from the extracellular side (cartoon mode). TMs and ECLs are colored slate. Compound 16 is shown as cyan sticks. C161^4.64^ and C173^5.32^ are shown as sticks and colored slate. (D) & (E) Interaction between compound 16 and MRGPRX1 from two views. The key residues are shown as sticks and colored slate. The polar interactions are shown as yellow dashed lines. (F) &(G) Bioluminescence resonance energy transfer assay (BRET) validation of the compound 16 binding pocket. Data are mean ± s.e.m. of n = 4 biological replicates. Emax, maximum effect; WT, wild type.

The orthosteric pocket of MRGPRX1 accommodating compound 16 is composed of residues majorly located on TM3/4/5/6 (Figure 2C). C161^4.64^ and C173^5.32^ form a disulfide bond, which is conserved among MRGPRXs[14] (Supplementary Figure 5). However, MRGPRX1 lacks the canonical disulfide bond between TM3 and ECL2 in other class A family GPCRs[28]. The disulfide bond substitution helps to reorganize the extracellular loops and maintain the wide-open orthosteric pocket.

### The interaction between compound 16 and MRGPRX1

Compound 16 adopts a hairpin conformation in the pocket due to the intramolecular π-π interaction of the 1-aminoisoquinoline and phenylmethyl groups (Figure 1A). The amino group of 1-aminoisoquinoline forms strong salt bridges to E157^4.60^ and D177^5.36^ (Figures 2D and E), which are two conserved residues in all MRGPRXs (Supplementary Figure 5). Alanine substitutions of E157^4.60^ and D177^5.36^ impair receptor activation in both BRET and calcium imaging assay, suggesting that these two acidic residues play a crucial role in ligand recognition (Figure 2F, Supplementary Figures 6A–C, Supplementary Table 1). Additionally, the variable residue Y99^3.29^ recognizes compound 16 through two types of interactions, the π-π interaction with the middle aromatic ring of compound 16 and polar interaction with the amino group of 1-aminoisoquinoline (Figures 2D and E). The Y99^3.29^A mutation abolishes the G_q_ coupling activity (Figure 2F, Supplementary Table 1). Thus, Y99^3.29^ is also essential for the recognition of compound 16. The residue L240^6.59^ contributes to compound 16 binding through hydrophobic interaction with the N-heterocyclic group of 1-aminoisoquinoline (Figures 2D and E). The potency of compound 16 to the mutant L240^6.59^A of MRGPRX1 is partially reduced in BRET assay (Figure 2G, Supplementary Table 1), whereas is not affected in the calcium imaging assay (Supplementary Figure 6D). The discrepancy may indicate that L240^6.59^ is less important for the compound 16-binding pocket. The corresponding residue L247^6.59^ in MRGPRX2 was reported to be critical for its agonist (R)-ZINC-3573 binding (Supplementary Figure 7A), and Y240 is located on ECL3 of MRGPRX4, which is important for the recognition of the agonist MS47134[14, 15] (Supplementary Figure 7B). Besides, the positive charge residue K96^3.26^ in MRGPRX1 is substituted by a serine in MRGPRX2 (Supplementary Figure 5). K96^3.26^ is close to compound 16 (Figures 2D and E), but S103^3.26^ is far away from (R)-ZINC-3573 (Supplementary Figure 7A). However, the alanine substitution of K96^3.26^ slightly affects the receptor activation (Figure 2G, Supplementary Table 1, Supplementary Figure 6E), suggesting that K96^3.26^ does not directly participate compound 16 recognition. In contrast, the K96^3.26^ in MRGPRX4 is critical for MS47134 interaction[14] (Supplementary Figure 7B). The variation of pocket residues may partly explain the agonist selectivity among MRGPRXs[12].

### Activation of MRGPRX1

The compound 16-MRGPRX1-G_αq_ complex structure exhibited TM rearrangement in the cytoplasmic half. The cytoplasmic ends of TM3 and TM6 are about 16 angstrom apart, consistent with other class A G protein-engaged GPCRs in an active conformation[28] (Figure 3A). Additionally, the conserved NP^7.50^XXY motif, which does not interact directly with G proteins but is essential for receptor activation[22], is conserved in MRGPRX1 (Figure 3B). However, MRGPRX1 shows significant differences in some classic motifs for receptor activation. Firstly, the conserved D(E)^3.49^R^3.50^Y^3.51^ motif on TM3 of most class A GPCRs forms an ionic lock in an inactive conformation and is broken upon activation[28]. In MRGPRX1, Y^3.51^ is replaced by C121^3.51^, E119^3.49^ interacts with R134^ICL2^ via a strong salt bridge, and R120^3.50^ interacts with T217^6.36^ to limit the movement of TM6 and Y359 on G_αq_ to stabilize the complex via polar interactions (Figure 3C). Secondly, the conserved motif P^5.50^I^3.40^F^6.44^ is substituted by an L187^5.46^L191^5.50^L110^3.40^F225^6.44^ motif, which constrains the conformation of TM3/5/6 (Figure 3D). Notably, the conserved W^6.48^ in the toggle switch of other GPCRs is replaced by G229^6.48^ in MRGPRX1. This vital substitution results in an inward movement of the extracellular half of TM6, narrowing the gap between TM3 and TM6 and forming a shallow orthosteric pocket (Figure 3E). The above structure features are highly conserved among MRGPRXs, suggesting the common activation mechanism despite distinct agonist recognition features[14, 15].

**Figure 3.**
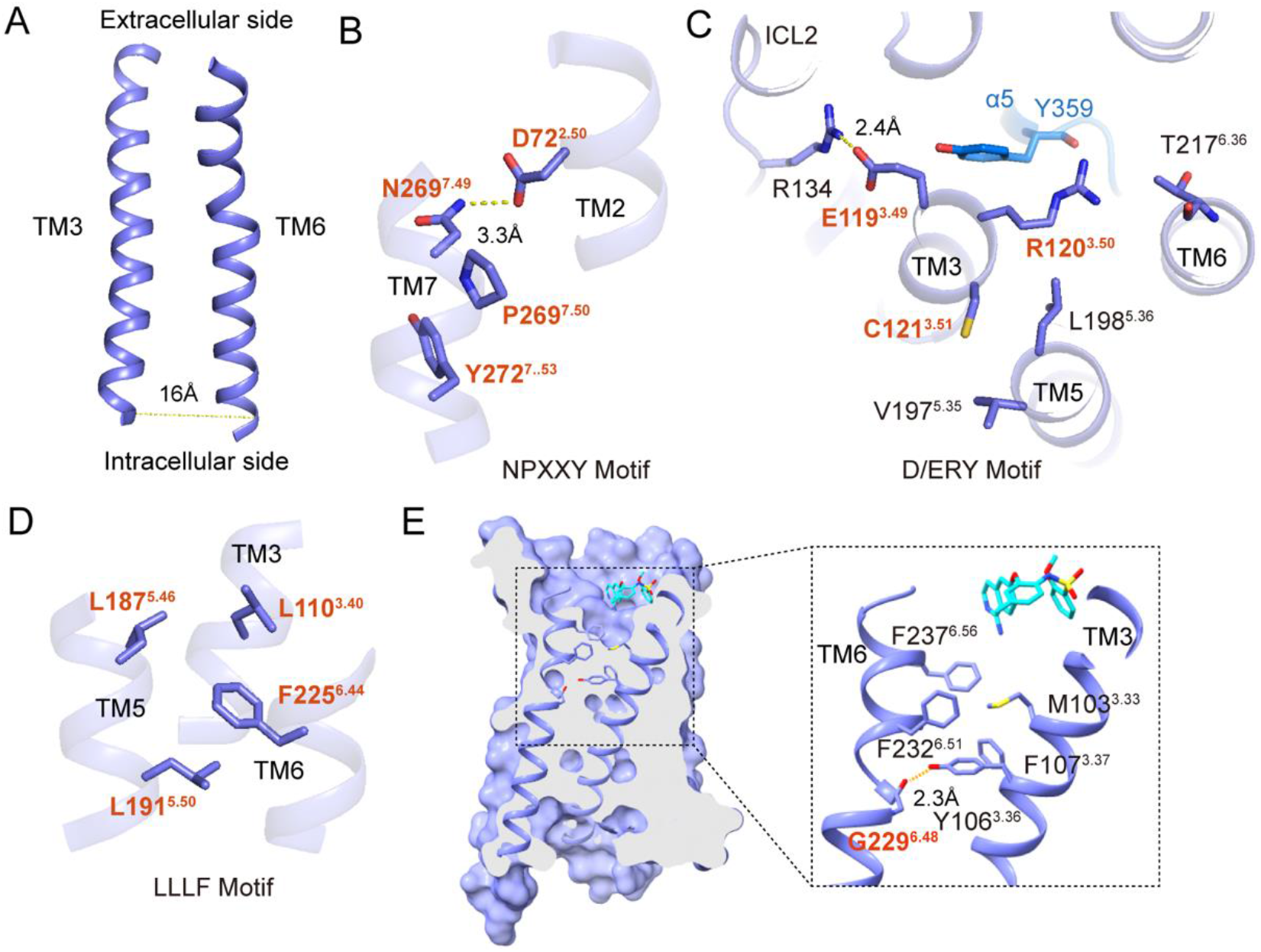
The conformational switches of MRGPRX1 in active state. (A) Structural representation of TMs arrangement between TM3 and TM6. The distance between the C-terminus of TM3 and TM6 is shown as a dashed line. (B) Magnified view of NPXXY motif. (C) Interactions between D/ERY motif and other residues in ICL2, TM3, TM5, TM6 and α5. Polar interactions are shown as yellow dashed lines. (D) Magnified view of LLLF motif. (E) Structural representation of the interactions near the substituted toggle switch G229^6.48^.

### The coupling of G_q_ to MRGPRX1

The coupling of G_q_ to MRGPRX1 is mainly maintained by interacting with residues on TM3, TM5, TM6, and ICL2 (Figure 4A). In most class A GPCRs, ICL2 does not interact with α5 of G protein directly. However, ICL2 plays a critical role in the G_q_-coupled MRGPRX1 activation process. E358 in α5 of G_αq_ interacts with R131^ICL2^ in MRGPRX1 via a strong salt bridge. N355 in α5 of G_αq_ forms a hydrogen bond with the carbonyl oxygen of P127^ICL2^ in MRGPRX1 (Figure 4B). Moreover, hydrophobic interactions are found between residues on α5 and TM3, TM5, and TM6, including V124^3.54^, I202^5.61^, L211^6.30^, and L214^6.33^ on the receptor and L352, L356, and L361 on the G_q_ (Figure 4C). Besides, R134^ICL2^ forms a hydrogen bond with R32 in αN of G_αq_ (Figure 4D). Similar G protein interfaces are observed in the complex structures of MRGPRX2 and MRGPRX4 previously reported (Supplementary Figures 8A and B).

**Figure 4.**
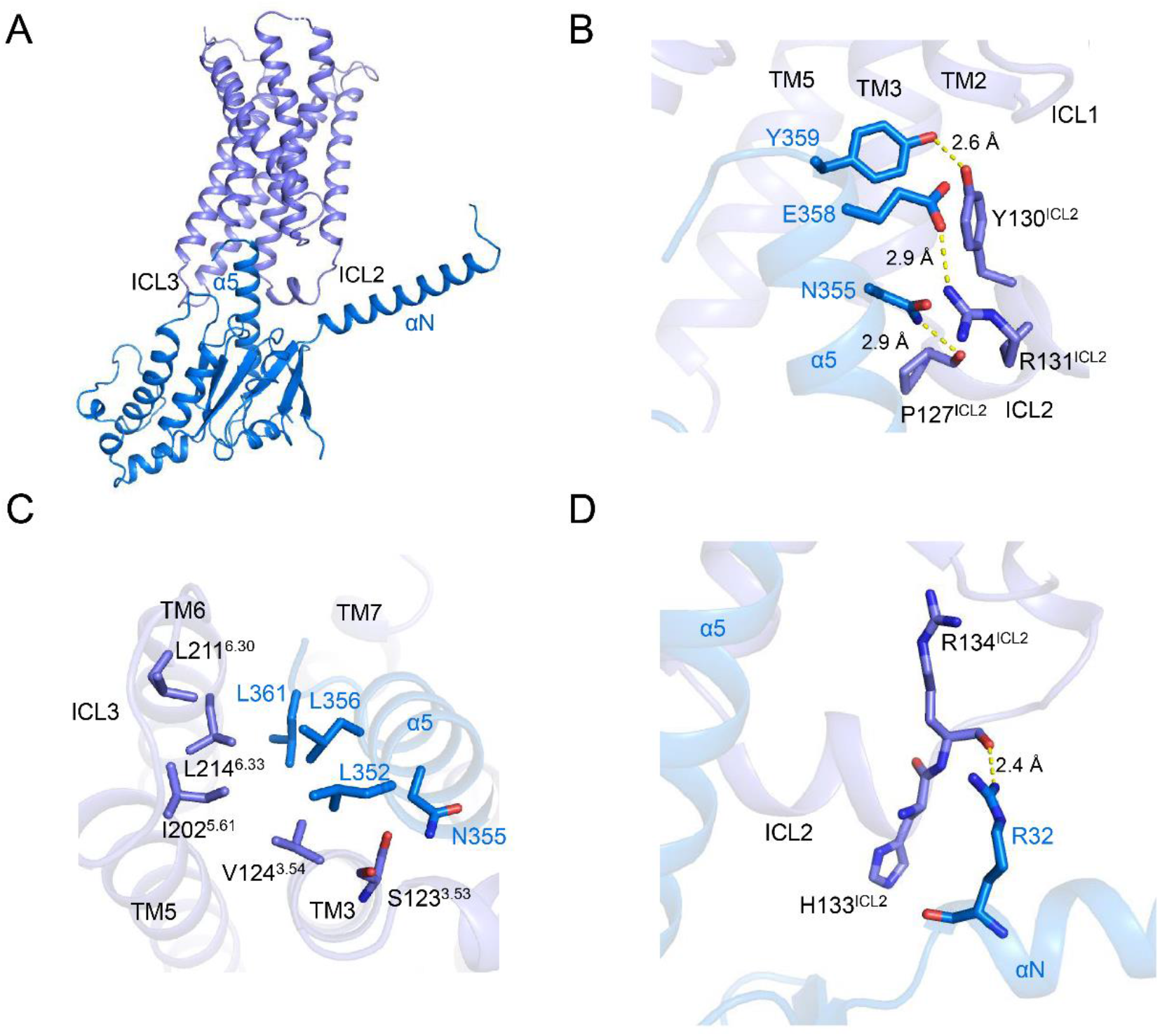
The coupling of MRGPRX1 to G_αq_. (A) Structural model shows the interaction between receptor and G_αq_ from the overall view. (B) Magnified view of ICL2-α5 helix interface. The polar interactions are shown as yellow dashed lines. (C) Hydrophobic interactions between TMs and the α5-helix of G_αq_. (D) Magnified view of ICL2-αN helix interface. The polar interaction is shown as yellow dashed line.

## Discussion

In this study, we used the NanoBiT strategy to determine the structure of compound 16-bound MRGPRX1 in complex with G_αq_ via cryo-EM. The structure reveals the common feature of shallow, broad, and wide-open orthosteric pockets in all MRGPRX members. We speculated that this shallow and broad pocket might easily accommodate various small compounds with distinct scaffolds. The less selectivity helps the receptor expand the ligand spectrum of itch sensation and facilitates the body’s quick response to the diverse exogenous stimulus.

Notably, the binding site of compound 16 is closed to TM3 and TM4 of MRGPRX1, while the binding site of (R)-ZINC-3573 is closed to TM5 and TM6 of MRGPRX2 (Figure 2C, Supplementary Figure 9A). The conformation of ECL2 in MRGPRX2 may prevent the ligand access to the corresponding position in MRGPRX1 (Supplementary Figure 9B). Similarly, the binding site of MS47134 is closed to TM2 and TM3 of MRGPRX4 (Supplementary Figure 9C). The inward movement of TM4, TM3, and ECL2 in MRGPRX4 may prevent the ligand access to the corresponding position in MRGPRX1 (Supplementary Figure 9D). Due to the distinct binding regions and the orthosteric pocket differences of these receptors, we evaluated the activation of compound 16 on MRGPRXs. As a result, MRGPRX1 is the only receptor that can be activated by compound 16 (Figure 5 A and Supplementary Figure 10). It further confirms that the MRGPRXs are differ in ligand recognition. These structural differences would give us clues to design agonists with improved specificity and potency.

**Figure 5.**
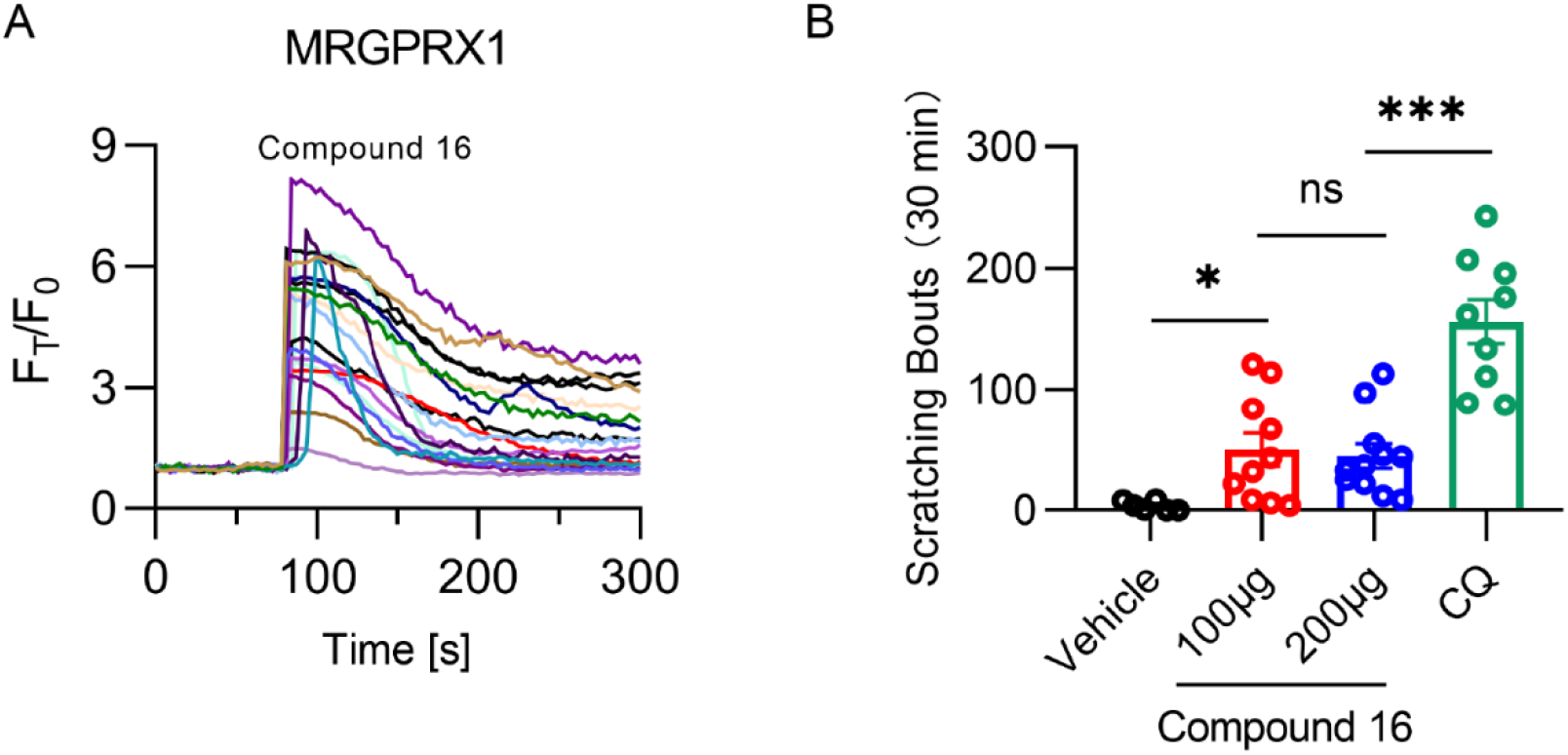
High selectivity of compound 16 to MRGPRX1 and its low itch risk. (A) Activation effect of compound 16 on MRGPRX1. Representative calcium traces of MRGPRX1 respond to compound 16 (500 nM). (B) Itch behavior evaluation of compound 16. Scratching responses induced by subcutaneous injection of vehicle (5% DMSO +95% saline containing 20% SBE-β-CD, n = 6, 3.667 ± 1.647), compound 16 (100 μg, n = 10, 50.20 ± 13.93; 200 μg, n = 11, 44.82 ± 9.978) and chloroquine (CQ, 200 μg, n = 9, 156.1 ± 18.24) in wild-type (WT) mice. Each dot represents an individual mouse. All data are presented as the mean ± SEM. ns, not significant, P > 0.5; *, P < 0.05; ***, P < 0.001.

Moreover, MRGPRX1 is reportedly involved in itch sensation[9] and pain inhibition[10]. Chloroquine (CQ) is a drug widely used in malaria treatment but results in the unbearable itch[29–31]. It is recently reported that MRGPRX1 mediates CQ-induced itch in humans[31]. Compound 16 was designed to inhibit chronic pain by targeting MRGPRX1. It is more abundant in the spinal cord than in the circulatory system, suggesting the lower risk of side effects caused by unexpected activation of MRGPRX1[16]. Given its high potency, high selectivity, and restricted distribution, compound 16 is a viable candidate drug worthy of more attention and further study[16]. We would evaluate if compound 16 also induces itch to limit its application. Here we used the scratching responses on the mouse model to evaluate the itch severity of compound 16 and chloroquine. As shown, compound 16 induces much less itch than a similar quantity of chloroquine (200 μg) (Figure 5B). In another word, compound 16 mainly participates in pain inhibition through MRGPRX1 activation. It provides a hint for the non-opioid pain-relieving molecule development. However, the molecular basis of chloroquine-mediated MRGPRX1 activation is unclear. Furthermore, studies on the downstream effectors of MRGPRX1, especially transient receptor potential vanilloid 1 (TRPV1) and transient receptor potential ankyrin 1 (TRPA1), show some conflicts. TRPV1 is usually regarded as an ion channels involved in pain sensation[32]. Sarah *et al*. found that TRPA1 is required for CQ-mediated itch[33]. Nevertheless, the tick salivary peptide IPDef1, an ancient invertebrate defensin peptide isolated from tick salivary, has been reported to evoke itch via MRGPRX1 and result in the activation of the downstream ion channel TRPV1 rather than TRPA1[34]. These data indicate that itch-sensing pathways and pain-sensing pathways appears to be indistinguishable. Thus, the differences in downstream signaling of MRGPRX1-mediated pain and itch sensations need further investigation.

## Materials and Methods

Detailed experimental procedures used in this study, including molecular cloning, cell culture, recombinant protein expression and purification, cryo-EM data collection, structure determination, and BRET and calcium imaging assays are available as SI Appendix and Methods.

## Data availability

Cryo-EM maps and atomic models have been deposited in the Electron Microscopy Data Bank under accession codes: EMD-34833 and in the Protein Data Bank under accession codes: 8HJ5. The raw data of BRET assay is shown in Supplementary Figure 11.

## Author contributions

B.G. designed and cloned the MRGPRX1 constructs for insect cell expression and biochemical assays, expressed and purified the complex for cryo-EM sample preparation, analyzed the data, prepared the figures and wrote the manuscript. L.Y.Y. processed the cryo-EM data, built the protein atomic model, refined the structures, analyzed the data, prepared the figures and assisted in the manuscript preparation. H.F.Y. accomplished the itch behavioral assay, performed calcium imaging assay, analyzed the data and prepared the figures. H.Z.J. prepared the grids and collected cryo-EM data. Y.A.C. assisted in the itch behavioral assay and calcium imaging assay. C.W. and R.L. assisted in the protein expression and purification. H.L.H. assisted in the structure validation. Z.J.C supervised the functional work and revised the manuscript. R.B.R. designed and supervised the whole project, participated in all data analysis and interpretation, wrote and revised the manuscript.

## Acknowledgments

We are grateful to Prof. H. Eric Xu from the Shanghai Institute of Materia Medica (SIMM) for donating the engineered G_αq_ chimera plasmid and Prof. Sheng Wang from the Shanghai Institute of Biochemistry and Cell Biology for donating the plasmids for functional assays. We thank Kobilka cryo-EM Center at the Chinese University of Hong Kong, Shenzhen, for supporting EM data collection. We also thank Ke Qiao from the core facility of Institute of Metabolic & Integrative Biology (IMIB) at Fudan University for her assistance with the user training. This work was supported by funds from National Natural Science Foundation of China project 32070525 (to Z.J. C.) and Shenzhen Science and Technology Program project JCYJ20220530140800001 (to Z.J. C.). B. G. were supported by Ganghong Youth Scholarship at the Chinese University of Hong Kong, Shenzhen.

## Methods

### Construct design

The full-length wild-type human MRGPRX1 (residues M1-S382) was cloned into pFastBac vector with a HA signal peptide, an N-terminal Flag tag, and a C-terminal 10xHis tag. A BRIL[21] protein was fused at the N-terminus of MRGPRX1 to improve the expression of the receptor. LgBit[22] was fused at the C-terminus of MRGPRX1 to stabilize the whole MRGPRX1-G_αq_ complex. There was no more modification for the MRGPRX1 sequence. Prof. H. Eric Xu from Shanghai Institute of Materia Medica donated the engineered G_αq_ chimera plasmid. This engineered G_αq_[24, 35] was designed based on a mini-G_αs/q_[23] skeleton with its N-terminus replaced by G_i1_ N-terminus for the binding of two G protein–stabilizing antibodies Nb35[26] and scFv16[27]. The wild-type G_β1γ2_ with HiBit[22] fused at the C-terminus of the β1 subunit was cloned into pFastBac-Dual vector. Ric8A is a molecular chaperone that has been reported to be essential for the biogenesis and signaling of G_αq_ subunits, and the involved functions include facilitating the proper folding of G_αq_ and promoting the formation of Gα guanine nucleotide binding pocket[36–38]. The full-length Ric8A was cloned into pFastBac vector. For Biochemical assay, the full-length wild-type MRGPRX1 and the MRGPRX1 point mutants were cloned into mammalian expression vector pCDNA 3.1 with an N-terminal Flag tag. All these constructs were generated with a standard PCR-based strategy and homologous recombination (CloneExpress One Step Cloning Kit, Vazyme).

### Expression and purification of scFv16

The scFv16 with a C-terminal 8×His tag was expressed in *Trichoplusia ni* Hi5 insect cells and purified precisely as previously described[27, 39, 40]. Briefly, the Hi5 insect cells (3.0-4.0 million cells per ml) were infected with the scFv16 virus produced by the Bac-to-Bac system (Invitrogen) for 96h. Then the medium was collected and pH balanced to pH 8.0 by adding Tris-base powder. Chelating agents were quenched by adding 1 mM nickel chloride and 5 mM calcium chloride. After incubation at room temperature (25 °C) for 1h by stirring constantly, the supernatant was isolated by centrifugation (4,750 rpm, 30 minutes, 4 °C) and incubated with Ni-sepharose (GE Healthcare) for 1h at room temperature with constant stirring. The resin was collected and then washed by washing buffer (20 mM HEPES pH7.5, 100 mM NaCl and 20 mM Imidazole). The protein was eluted by elution buffer (20 mM HEPES pH 7.5, 100 mM NaCl and 250 mM Imidazole) and then incubated with HRV-3C protease at 4 °C for 2h to remove the C-terminal 8×His tag. The cleaved protein was further loaded onto Superdex 200 Increase 10/300 GL column (GE healthcare) with running buffer (20 mM Hepes pH 7.5, 100 mM NaCl). ScFv16 peak fraction was collected, flash-frozen, and stored at −80°C until use.

### Expression and purification of Nb35

Nanobody-35 (Nb35) was expressed in the *Escherichia coli* BL21 (DE3) and purified as previously reported[26, 35]. Nb35 was cultured in LB with 50 μg/mL ampicillin at 37°C, 220 rpm for about 3h until OD_600_ reached 1.0. Then IPTG was added to induce the protein expression at 25°C, 220 rpm with a final concentration 1 mM. Then the *E. coli* bacteria were collected by centrifugation (4,000 rpm, 20 minutes, 4 °C) after 16h. The pellets were resuspended in lysis buffer (25 mM Hepes pH 7.5, 150 mM NaCl, 1mM PMSF) and disrupted by ultrasonication. The supernatant was isolated by centrifugation (14,000 rpm, 60 minutes, 4 °C) and incubated with Ni-NTA resin (Qiagen) for 1h. The resin was washed by washing buffer (25 mM HEPES pH 7.5, 150 mM NaCl and 25 mM Imidazole and then eluted by elution buffer (25 mM HEPES pH 7.5, 150 mM NaCl and 250 mM Imidazole). The eluted protein was concentrated and loaded onto Superdex 200 Increase 10/300 GL column (GE healthcare) with running buffer (25 mM Hepes pH 7.5, 150 mM NaCl). The Nb35 monomeric fractions were pooled and stored at −80°C for further use.

### Expression and purification of MRGPRX1-G_αq_ complex

The complex was expressed in *Spodoptera frugiperda* (*Sf9*) insect cells. Baculovirus preparation was accomplished based on the baculovirus system manual (Invitrogen). For expression, the Sf9 cells were cultured in serum-free medium (UK1000, UNION-BIOTECH) to 2.0 million cells per mL and co-infected with BRIL-MRGPRX1-LgBit, engineered G_αq_, G_β1γ2_-HiBit, and Ric8A at a virus ratio 1:1:1:1 for 48h. The complex cell pellets were resuspended in lysis buffer (25 mM HEPES pH 7.5, 150 mM NaCl, 5% Glycerol, 10 mM MgCl_2_, 20 mM KCl, 5mM CaCl_2_) supplemented with Protease Inhibitors (1mM PMSF, 2 μg/mL Aprotinin, 5 μg/mL Leupeptin and 1 μg/mL Pepstatin). The suspension was homogenized and incubated with 10 μM compound 16 (MCE), scFv16 (20 μg/mL), Nb35 (15 μg/mL), and apyrase (25 mU/mL, NEB) for 1h at room temperature. Then 0.5% (w/v) lauryl maltose neopentylglycol (LMNG) (Anatrace) and 0.05% (w/v) CHS (Anatrace)) was added into the suspension to solubilize the membrane. After extracting for 2h at 4 °C, the supernatant was isolated by centrifugation (18,000 rpm, 40 minutes, 4 °C. The flag resin was washed with 10 column volumes washing buffer 1 (25 mM HEPES pH 7.5, 150 mM NaCl, 5% Glycerol, 5 mM MgCl_2_, 5mM CaCl_2_, 10 μM compound 16, 0.1% (w/v) LMNG and 0.01% (w/v) CHS), and then washed with 10 column volumes washing buffer 2 (25 mM HEPES pH 7.5, 150 mM NaCl, 5% Glycerol, 5 mM MgCl_2_, 5mM CaCl_2_, 10 μM compound 16, 0.01% (w/v) LMNG, and 0.001% (w/v) CHS). The complex was eluted with elution buffer (25 mM HEPES pH 7.5, 150 mM NaCl, 5% Glycerol, 5 mM MgCl_2_, 5mM CaCl_2_, 10 μM compound 16, 200 μg/mL Flag peptide (GenScript), 0.01% (w/v) LMNG, 0.005% GDN (Anatrace), and 0.001% (w/v) CHS). The final complex elution was concentrated to less than 2 mL using an Amicon Ultra Centrifugal Filter (MWCO 100 kDa) and incubated with 50 μM compound 16 for 1h at 4 °C. Then the complex was subjected to Superdex 200 Increase 10/300 GL column (GE healthcare) with running buffer (25 mM HEPES pH 7.5, 150 mM NaCl, 2 mM MgCl_2_, 10 μM compound 16, 0.00075% (w/v) LMNG, 0.00025% (w/v) GDN, 0.000075% (w/v) CHS) to remove uncoupled components. The monomeric peak fractions were collected and concentrated to the final concentration 12 mg/mL for cryo-EM grid preparation.

### Cryo-EM sample preparation and data acquisition

3.5 μL of 12 mg/mL the concentrated complex was applied to glow-discharged holey carbon-coated grids (Quantifoil 200 mesh, Au R1.2/1.3). The grids were blotted for 3.5s and flash-frozen in liquid ethane using a Vitrobot (Mark IV, Thermo Fisher Scientific). Images were recorded on a 300kV Titan Krios G3i electron microscope (Thermo Fisher Scientific) equipped with Gatan K3 Summit direct detector and a GIF Quantum energy filter (slit width 20 eV). Movie stacks were collected using SerialEM[41] in counting mode at a magnification of 105,000x with the corresponding pixel size of 0.85Å. Movies stacked with 50 frames were exposed for 2 s. Two data sets of the same sample were collected, including 3,447 and 2,799 movies separately. Two data sets were recorded at a total dose of about 56.15 e/Å2 and 58.32 e/Å2, respectively. The defocus range was set from −1.0 μm to −2.0 μm.

### Data Processing

Data processing was performed using cryoSPARC[42]. Movies frames were aligned using Patch motion. CTF estimation was performed using Patch CTF. Particles were first picked using a blob picker with partial micrographs. 2D templates were generated by 2D classification. Particle picking of all micrographs was performed by a template picker. 6,687,388 particles from 6,246 micrographs were extracted using a box size of 288 pixels and cropped into 72 pixels. After two rounds of 2D classification and one round of ab initio reconstruction, 1,808,631 particles were selected and re-extracted using a box size of 288 pixels. Ab initio reconstruction using partial particles was performed, and 279,237 particles were removed, remaining 1,529,394 particles. After that, one round of non-uniform and local refinement was performed. One round of ab initio reconstruction was performed again, generating a new dataset with 1,006,848 particles. Finally, one round of non-uniform and local refinement was performed, generating a 3.0 Å map.

### Model building and refinement

The initial complex model was built using the structure of MRGPRX2 (PDB code: 7S8N) and Nb35 (PDB code: 7F4H) as templates. Models are then fitted into the density map and manually adjusted and rebuilt in COOT[43]. The restraint files of compound 16 and CHS were generated by Phenix. elbow package[44]. The complete model was finally refined in Phenix using real-space refinement with secondary structure and geometry restraints[45] and COOT. Overfitting of the model was checked by refining the model using one of the two independent maps from gold-standard refinement and calculating FSC against both half maps[46]. The final model was validated using Molprobity[47](Table S1). Structural figures were prepared in PyMOL (https://pymol.org/2/), UCSF Chimera[48], and UCSF ChimeraX[49].The initial complex model was built using the structure of MRGPRX2 (PDB code: 7S8N) and Nb35 (PDB code: 7F4H) as templates. Models are then fitted into the density map and manually adjusted and rebuilt in COOT[43]. The restraint files of compound 16 and CHS were generated by Phenix. elbow package[44]. The complete model was finally refined in Phenix using real-space refinement with secondary structure and geometry restraints[45] and COOT. Overfitting of the model was checked by refining the model using one of the two independent maps from gold-standard refinement and calculating FSC against both half maps[46]. The final model was validated using Molprobity[47](Table S1). Structural figures were prepared in PyMOL (https://pymol.org/2/), UCSF Chimera[48], and UCSF ChimeraX[49].

### Bioluminescence resonance energy transfer assay (BRET)

BRET assays[50] were performed as previously reported[39] to measure MRGPRX1-mediated G_q_ protein activation. Briefly, HEK293T cells (ATCCCRL-11268; mycoplasma free) were co-transfected at a 1:1:1:1 ratio of receptor: G_αq_-Rluc8: G_β_: G_γ_-GFP2. After at least 18h, transfected cells were harvested and reseeded in opaque white bottom 96-well assay plates (Beyotime) at a density of 30,000-50,000 cells per well (DMEM plus 10% FBS). The medium was removed after 24h. Cells were incubated with 40 μL 7.5 μM coelenterazine 400a (Goldbio) in drug buffer (1×Hank’s balanced salt solution (HBSS), 20 mM HEPES, pH 7.4, 0.3%BSA) for 2 minutes, and then treated with 20 μL compounds prepared in drug buffer at serial concentration gradient for additional 5 minutes. Plates were read in an LB940 Mithras plate reader (Berthold Technologies) with 395-nm and 510-nm emission filters. BRET ratios were calculated as the ratio of GFP2 emission (510 nm) to Rluc8 emission (395 nm) and analyzed by GraphPad prism 8.0.

### Behavioral studies

Wild-type C57BL/6J mice (8-week-old males) in acute itch behavioral tests were purchased from the Disease Control and Prevention Center of Hubei Province in China. Vehicle and MRGPRX1 agonists were subcutaneously injected into the nape of neck of mice after acclimatization. Scratching behavior in mice was observed for 30 min following injection. A bout of scratching was defined as a scratching movement with the hind paw directed at the area of the injection site. Then, the scratching bouts directing at the injected site were quantified.

### Calcium imaging

HEK293T cells were loaded with 5 μM Fluo-4AM (Yeasen) for 30-45 min in the dark, supplemented with 0.01% Pluronic F-127 (wt/vol, Yeasen), in 1× Hank’s Balanced Salt Solution (HBSS) containing 140 mM NaCl, 5 mM KCl, 10 mM HEPES, 2 mM CaCl_2_, 2 mM MgCl_2_ and 10 mM d-(+)-glucose (pH 7.4). After washing 3 times with HBSS, emission at 520 nm was detected from 488 nm excitation for Fluo-4AM. Data were analyzed from at least three repeated experiments at 80-200 cells.

---

**Supplementary Figure 1.**
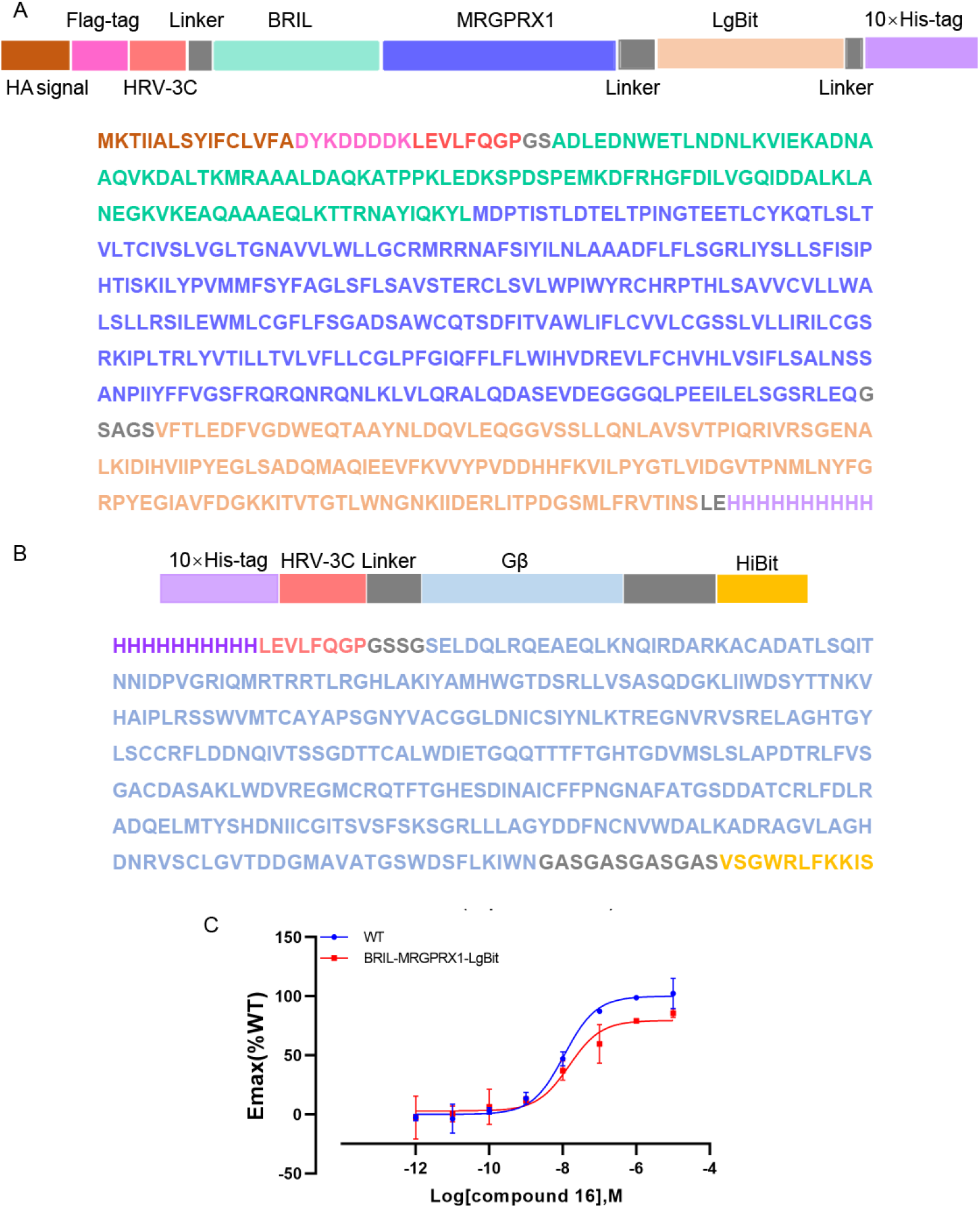
MRGPRX1 construct and G_β_-HiBit construct used in this study. (A) Schematic representation and amino acid sequence of MRGPRX1 construct. Haemagglutinin (HA) signal (reddish brown), Flag tag (pink), HRV-3C protease cleavage sites (deepsalmon), fusion protein BRIL (greencyan), S1PR1 (slate), LgBit (wheat), His tag (violet), and linker (gray). (B) Sequence of G_β_-HiBit construct. His tag (violet), HRV-3C protease cleavage sites (deepsalmon), G_β_ (light blue), HiBit (orange), and linker (gray). (C) Potency evaluation of the compound 16 induced G_αq_ dissociation in MRGPRX1-WT and BRIL-MRGPRX1-LgBit overexpressing cells. Data are mean ± s.e.m. of n = 4 biological replicates. Emax, maximum effect; WT, wild type.

**Supplementary Figure 2.**
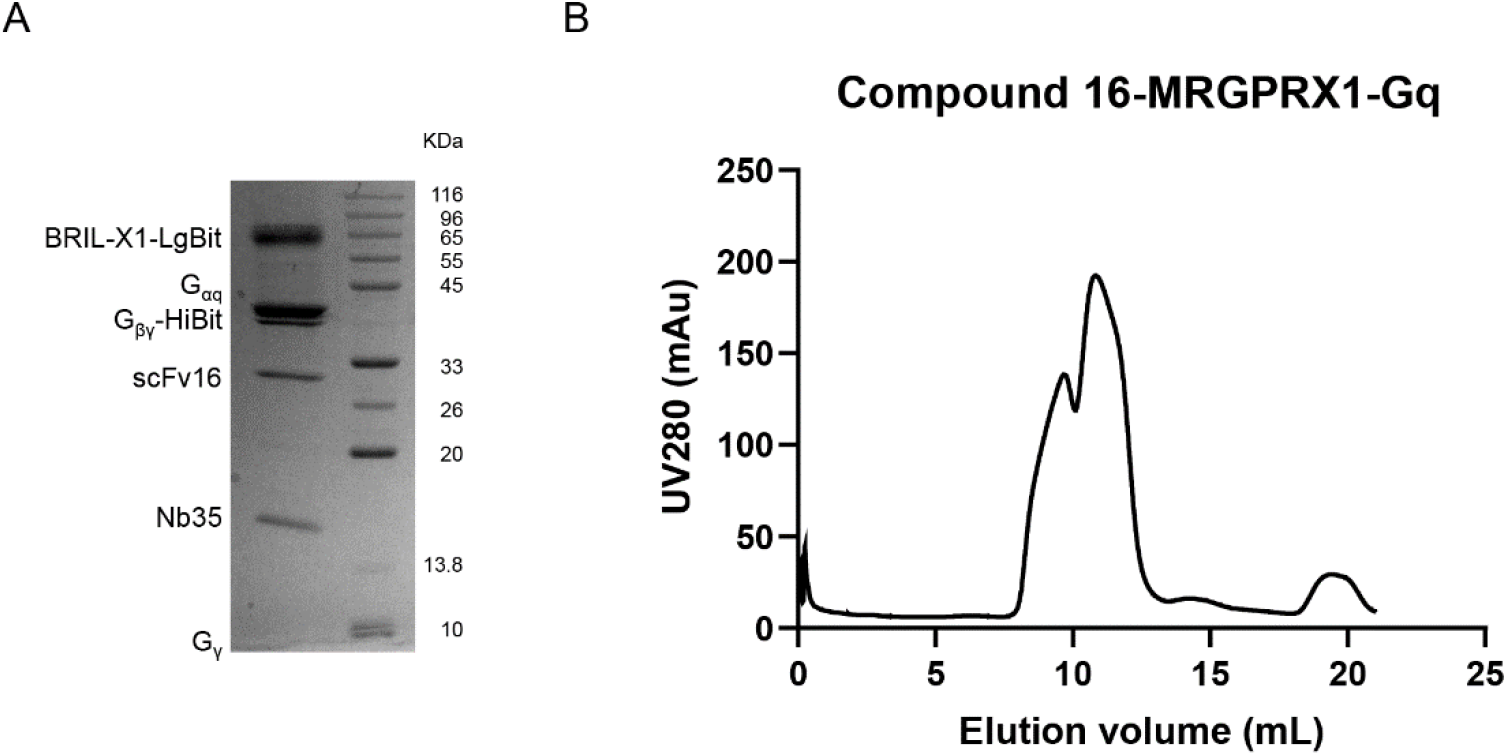
Compound 16-MRGPRX1-G_αq_s complex purification. (A) SDS-PAGE of the Flag-purified compound 16-MRGPRX1-G_αq_ complex. (B) Final size exclusion chromatography elution profile of the complex.

**Supplementary Figure 3.**
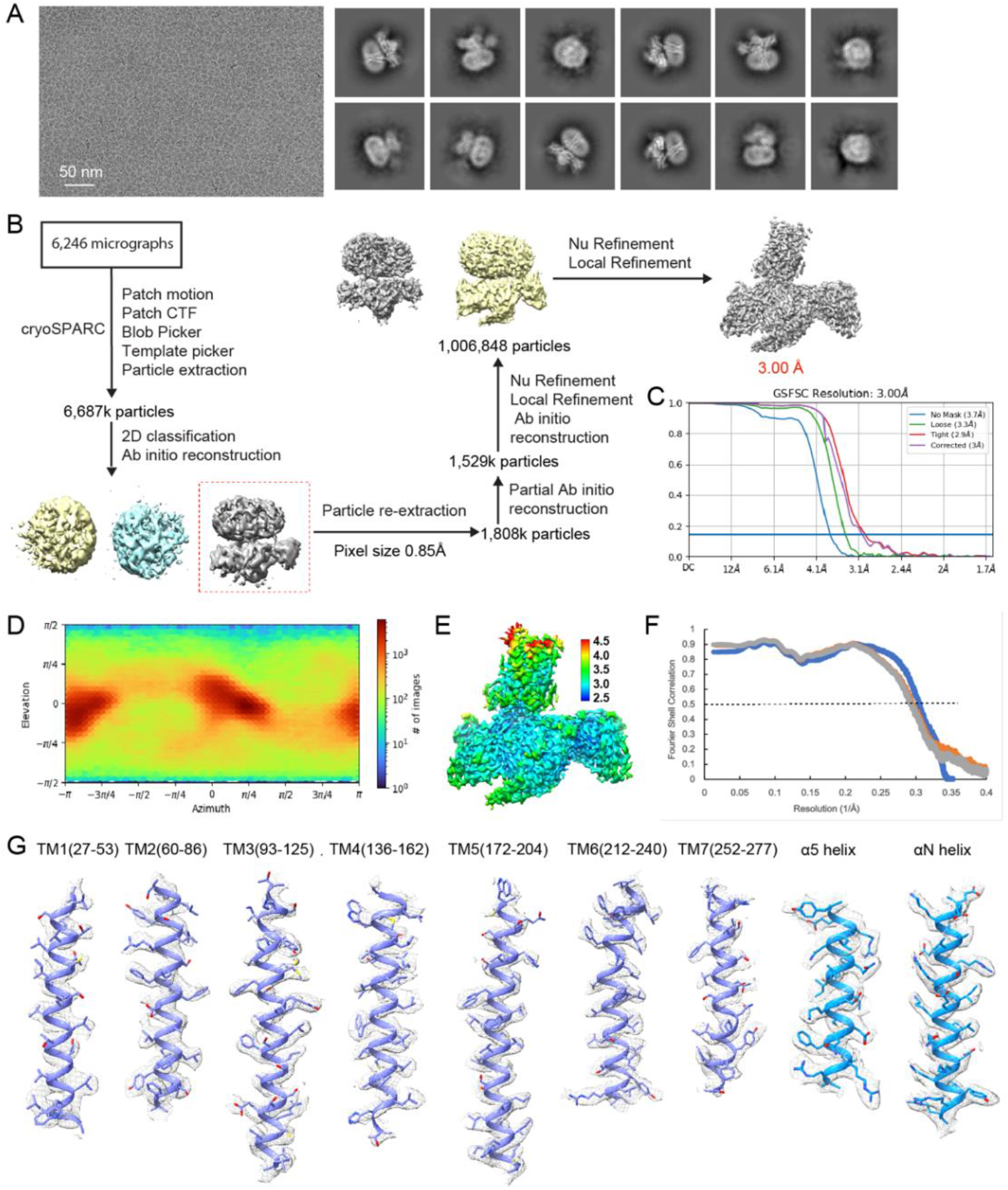
Structural determination of compound 16 bound MRGPRX1-G_q_ complex. (A) Representative micrograph and 2D classes. (B) Flowchart for cryo-EM data processing. Details can be found in the Methods. (C) Fourier shell correlation (FSC) curves of the final refined cryoEM map. (D) Angular distribution of the particles used for the final reconstructions. (E)Local resolution of the final map estimated by cryoSPARC. (F) Fourier shell correlation (FSC) between map and model. FSC curve of the final refined model against the full map, colored in blue. FSC curve of the model refined against the first half map against the same map, colored in orange. FSC curve of the model refined against the first half map against the second half map, colored in gray. (G) The density maps of the transmembrane helix of MRGPRX1, α5 helix and αN helix are shown as mesh.

**Supplementary Figure 4.**
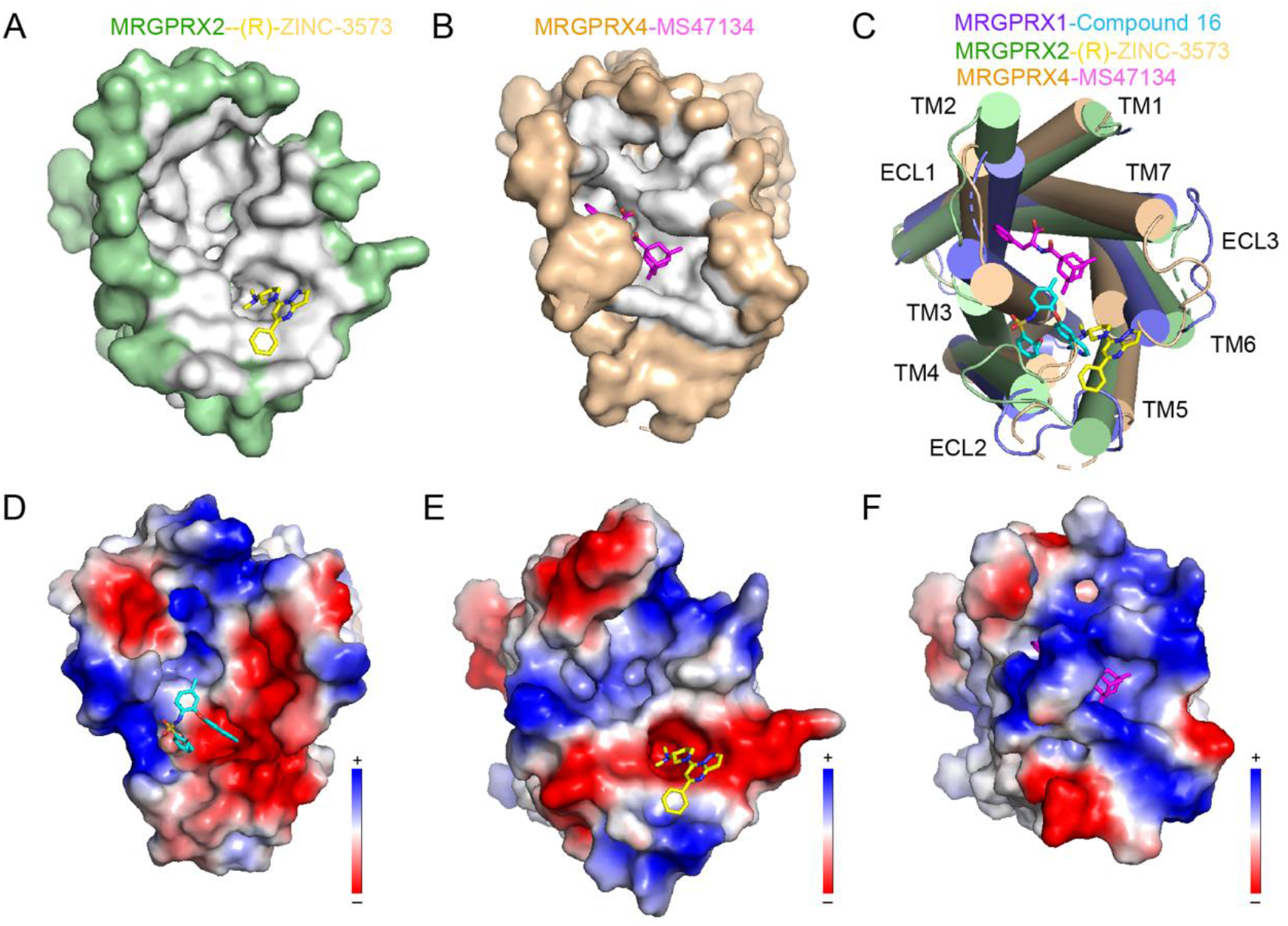
Ligand-binding pocket of G_q_-coupled MRGPRX1, MRGPRX2 and MRGPRX4. (A) Top view of the (R)-ZINC-3573-binding pocket from the extracellular side (surface mode). Pocket is colored gray, and (R)-ZINC-3573 is shown as yellow sticks. (B) Top view of the MS47134-binding pocket from the extracellular side (surface mode). Pocket is colored gray, and MS47134 is shown as magenta sticks. (C) Structural comparison of MRGPRX1-G_q_-compound 16 complex with MRGPRX2-G_q_-(R)-ZINC-3573 complex (PDB code: 7S8N) and MRGPRX4-G_q_-MS47134 complex (PDB code: 7S8P) in an extracellular view (cartoon mode). MRGPRX1, MRGPRX2, and MRGPRX4 are colored slate, palegreen, and wheat, respectively. Compound 16, (R)-ZINC-3573, and MS47134 are colored cyan, yellow, and magenta, respectively. (D) Electrostatic surface representation of the MRGPRX1 extracellular pocket calculated using PyMOL with compound 16 shown as cyan sticks. Red, negative; blue, positive. (E) Electrostatic surface representation of the MRGPRX2 extracellular pocket calculated using PyMOL with (R)-ZINC-3573 shown as yellow sticks. Red, negative; blue, positive. (F) Electrostatic surface representation of the MRGPRX2 extracellular pocket calculated using PyMOL with MS47134 shown as magenta sticks. Red, negative; blue, positive.

**Supplementary Figure 5.**
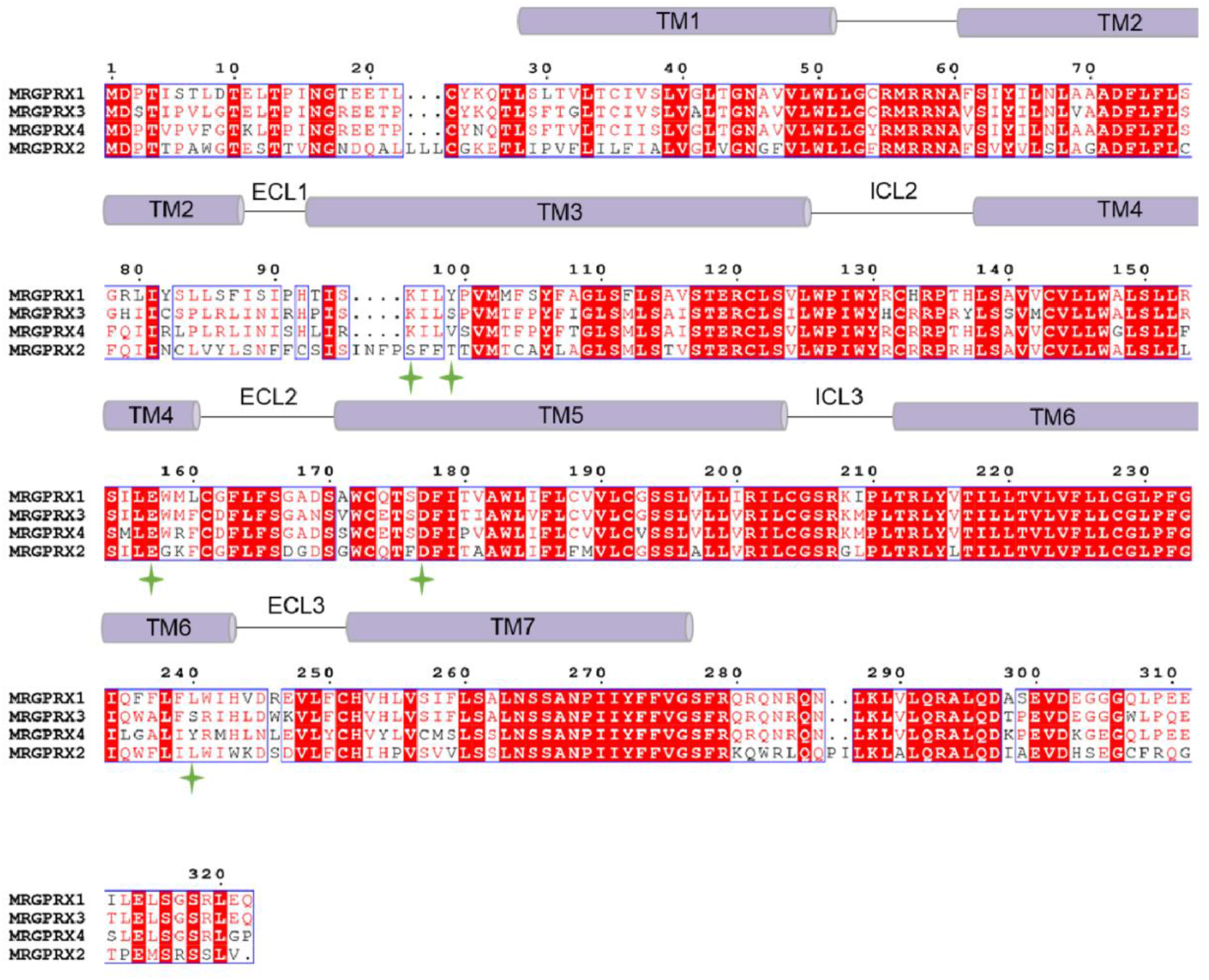
Sequence alignment of Human MRGPRX1, MRGPRX2, MRGPRX3 and MRGPRX4. The sequence alignment was created using Clustalw[51] and ESPript 3.0 servers[52]. TMs for MRGPRX1 receptor are shown as violet columns. The key ligand-binding pocket residues are shown as yellow green asterisk.

**Supplementary Figure 6.**
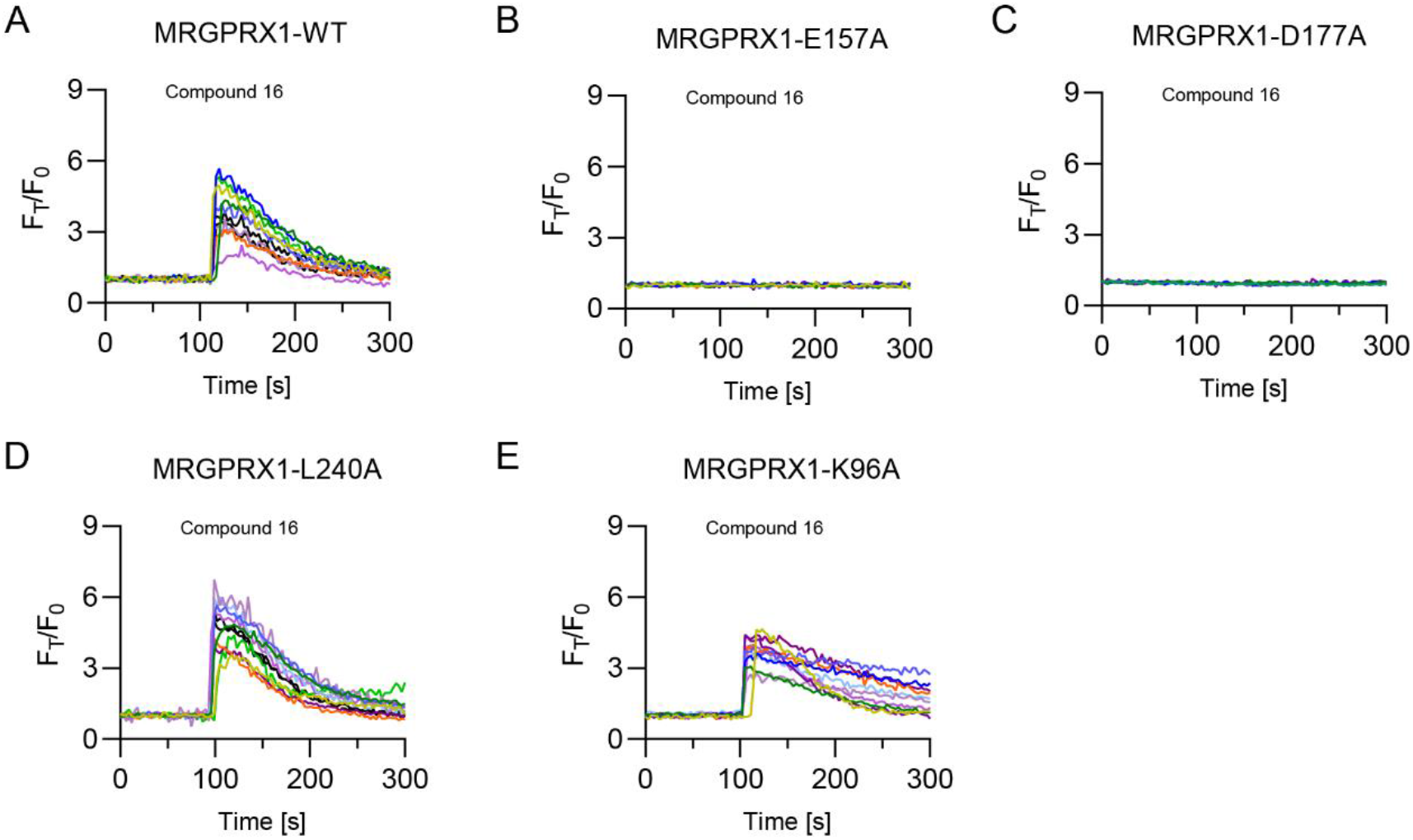
Calcium imaging validation of the compound 16 binding pocket. Representative calcium traces of MRGPRX1-WT (A), E157A (B), D177A (C), Y99A (D), L240A (E), and K96A (F) respond to compound 16 (500 nM).

**Supplementary Figure 7.**
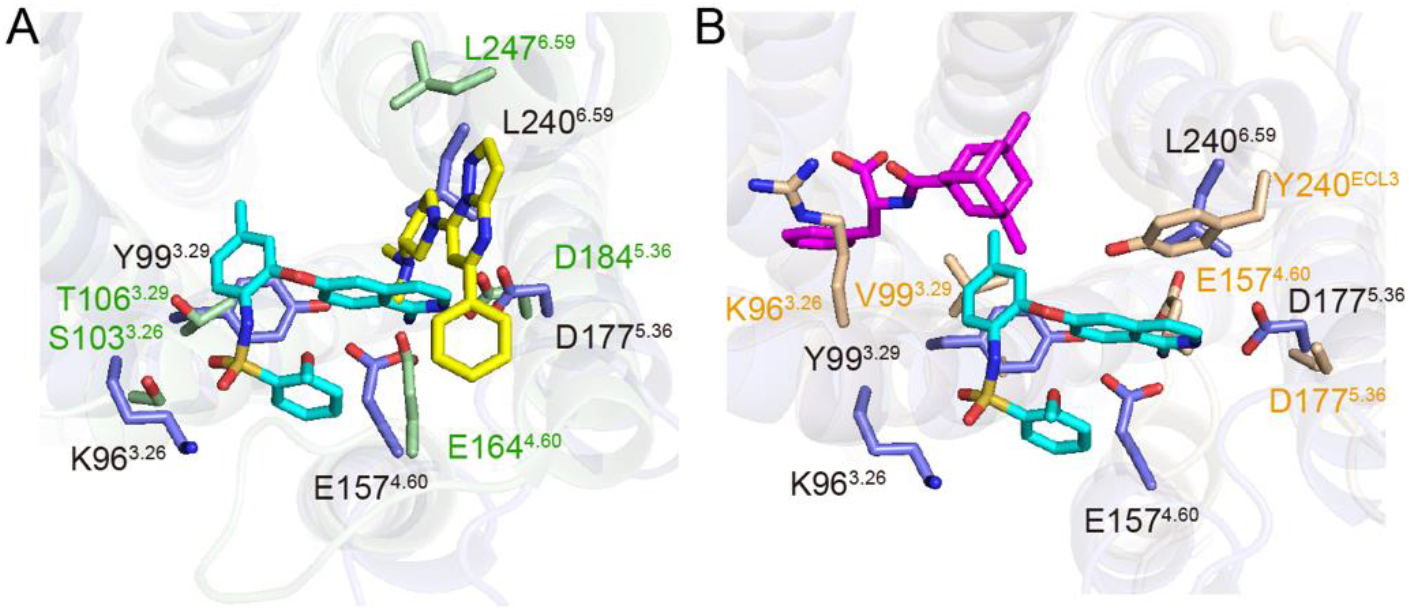
Detailed structure comparison of ligand-binding pocket among MRGPRX1, MRGPRX2 and MRGPRX4. (A) A comparison of the ligand-binding pocket between MRGPRX1 and MRGPRX2 in a magnifying view. The key residues in MRGPRX1 are colored slate, and the corresponding residues in MRGPRX2 are colored palegreen. Compound 16 and (R)-ZINC-3573 are colored cyan and yellow, respectively. (B) A comparison of the ligand-binding pocket between MRGPRX1 and MRGPRX4 in a magnifying view. The key residues in MRGPRX1 are colored slate, and the corresponding residues in MRGPRX2 are colored wheat. Compound 16 and MS47134 are colored cyan and magenta, respectively.

**Supplementary Figure 8.**
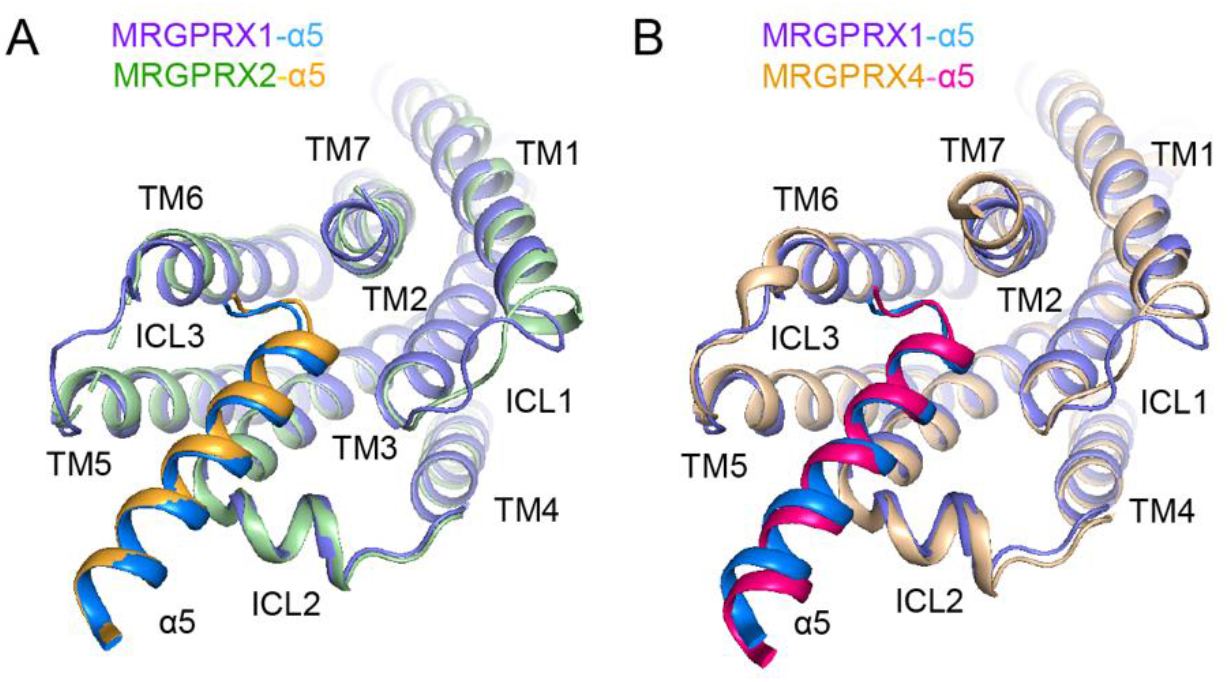
Comparison of the engagement of α5 helix in structure of MRGPRX1, MRGPRX2, and MRGPRX4. (A) A comparison of the engagement of α5 helix to receptor between the MRGPRX1 and MRGPRX2, viewing from the TM5, TM3, and ICL2 front angle. MRGPRX1, MRGPRX2, α5 helix in MRGPRX1, and α5 helix in MRGPRX2 are colored slate, palegreen, marine and orange, respectively. (B) A comparison of the engagement of α5 helix to receptor between the MRGPRX1 and MRGPRX4, viewing from the TM5, TM3, and ICL2 front angle. MRGPRX1, MRGPRX4, α5 helix in MRGPRX1, and α5 helix in MRGPRX4 are colored slate, wheat, marine and magenta, respectively.

**Supplementary Figure 9.**
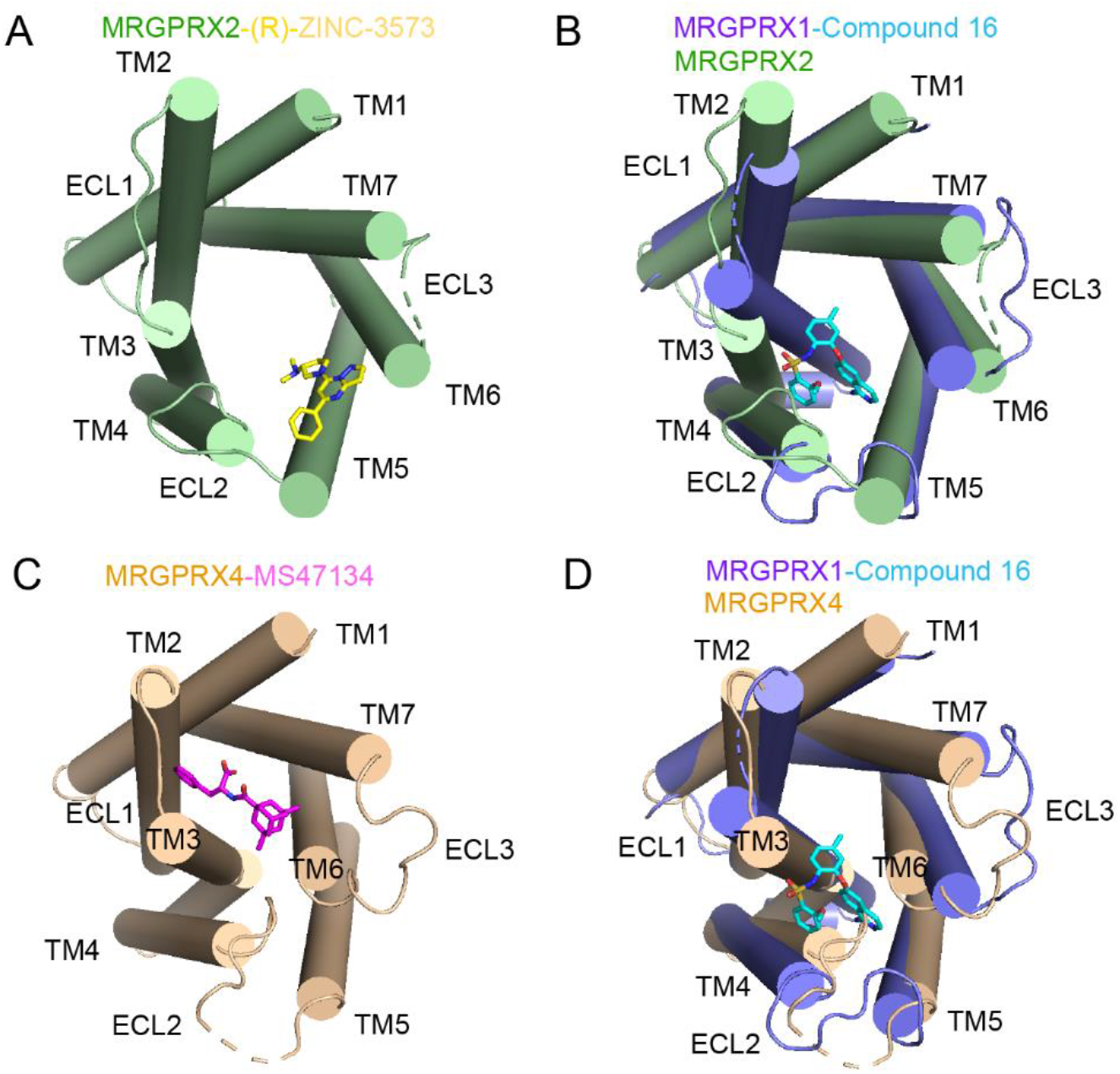
Structural models of human MRGPRX2 and MRGPRX4 in an extracellular view. (A) Top view of the (R)-ZINC-3573-binding pocket from the extracellular side (cartoon mode). MRGPRX2 is colored palegreen, and (R)-ZINC-3573 is shown as yellow sticks. (B) Top view of the superperimposed MRGPRX1-compound 16 with MRGPRX2 (cartoon mode). MRGPRX1 is colored slate, and compound 16 is shown as cyan sticks. MRGPRX2 is colored palegreen. (C) Top view of the MS47134-binding pocket from the extracellular side (cartoon mode). MRGPRX4 is colored wheat, and MS47134 is shown as magenta sticks. (D) Top view of the superperimposed MRGPRX1-compound 16 with MRGPRX4 (cartoon mode). MRGPRX1 is colored slate, and compound 16 is shown as cyan sticks. MRGPRX4 is colored wheat.

**Supplementary Figure 10.**
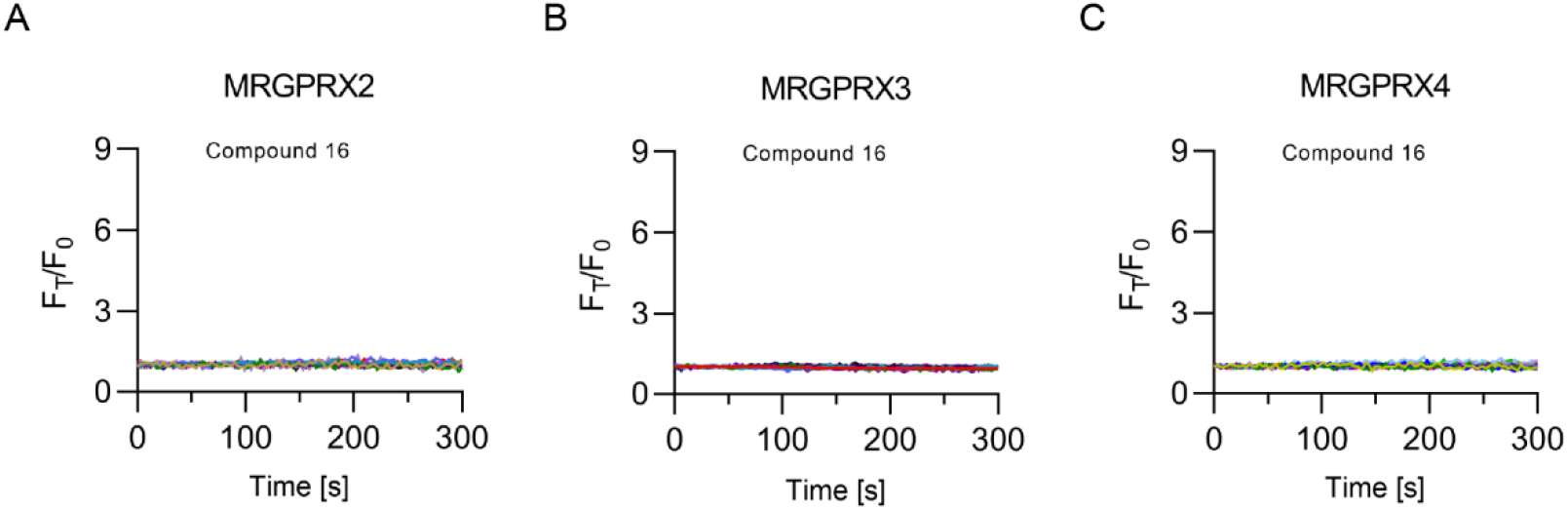
Activation effect of compound 16 on human MRGPRXs. Representative calcium traces of MRGPRX2 (A), MRGPRX3 (B), and MRGPRX4 (C) respond to compound 16 (500 nM). Details are described in methods.

**Supplementary Figure 11.**
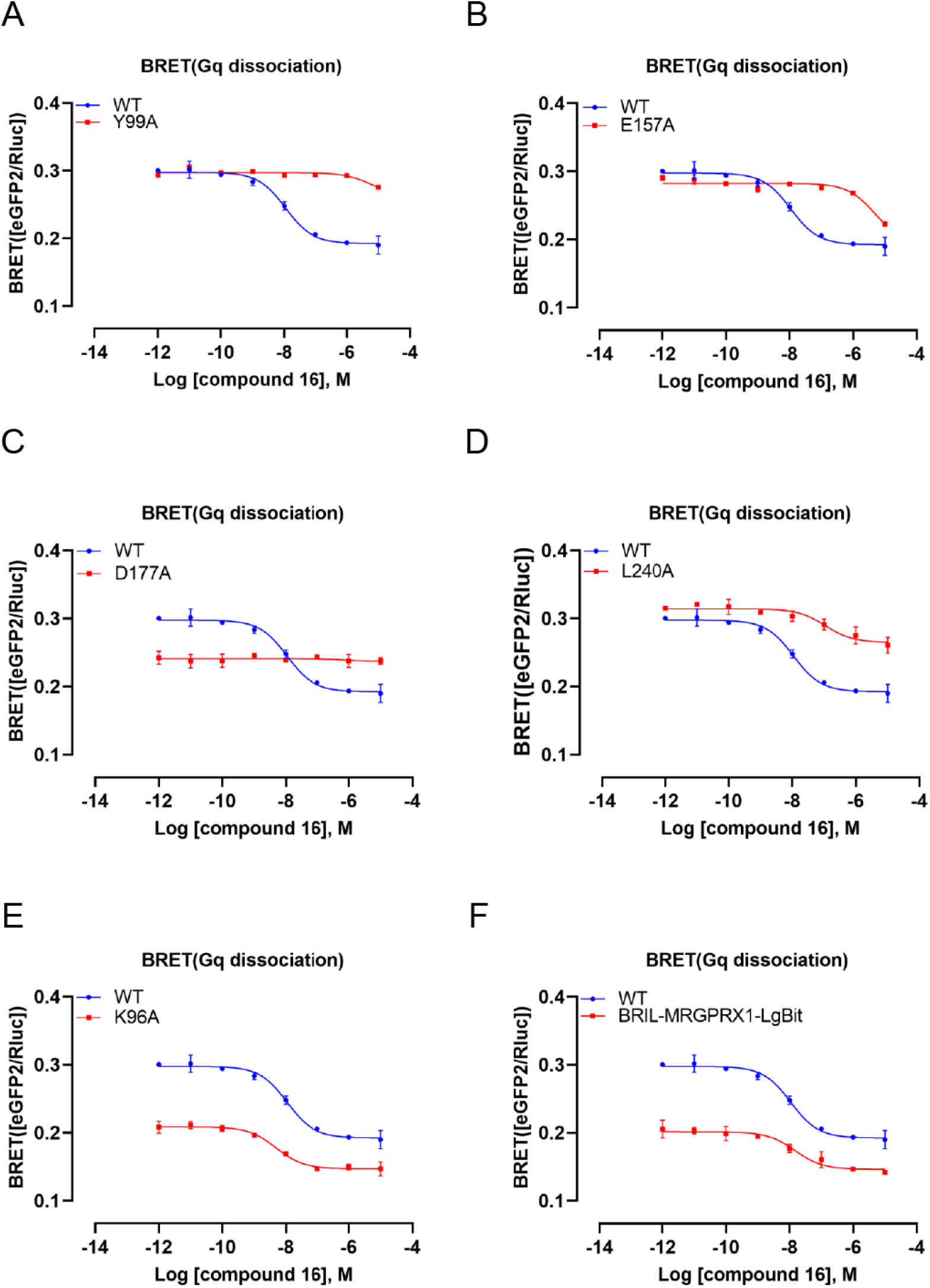
Raw data of the BRET assay. Dose response curves comparison of WT with Y99A (A), E157A (B), D177A (C), K96A(E), and BRIL-MRGPRX1-LgBit. Data are mean ± s.e.m. of n = 4 biological replicates. WT, wild type.

**Supplementary Table 1.**
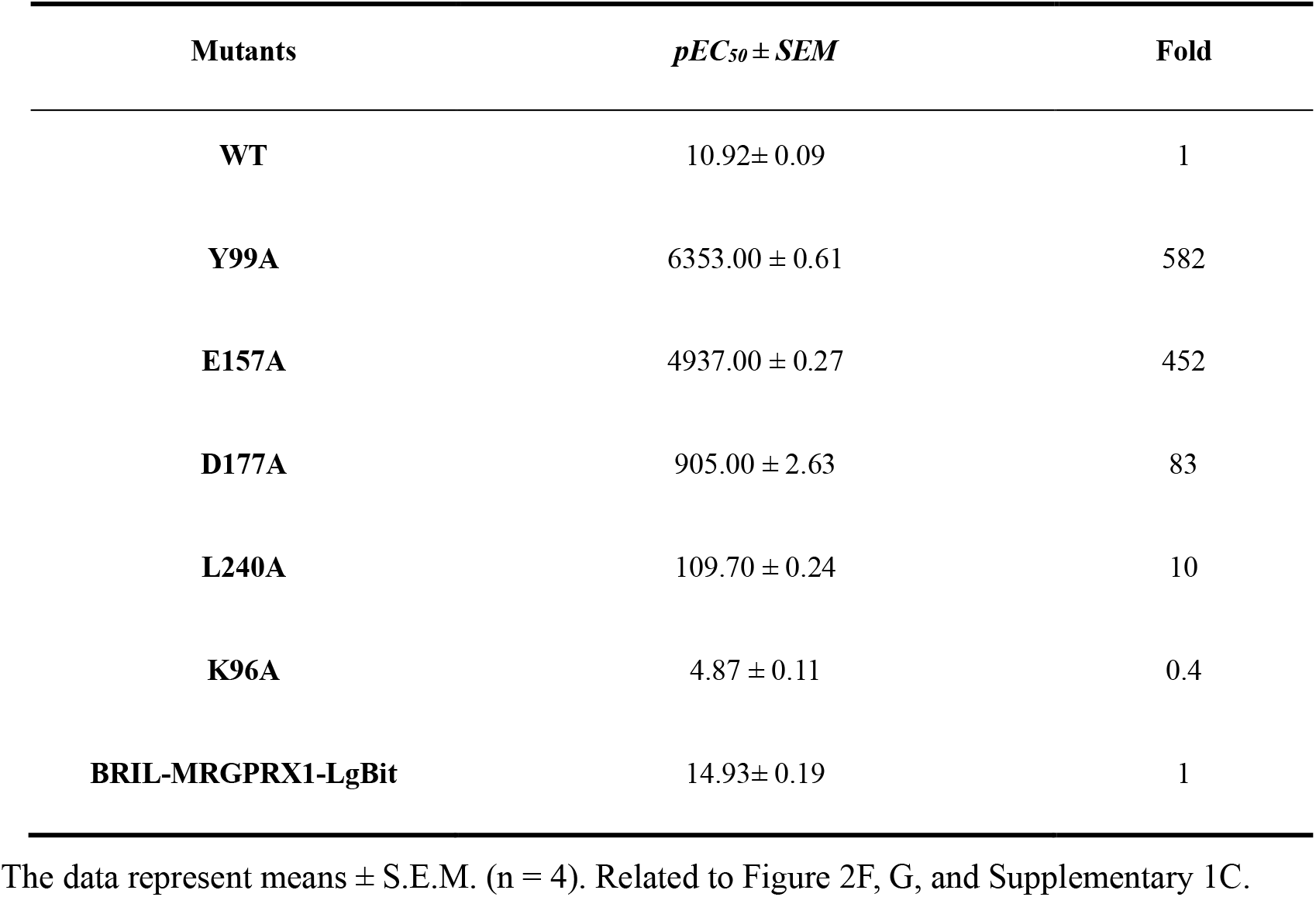
The potencies of Compound 16 with the wild type MRGPRX1 receptor or its mutants.

**Supplementary Table 2.** Cryo-EM data collection, refinement and validation statistics.

|  | compound 16 bound<br>(EMDB-34833)<br>(PDB 8HJ5) |
| --- | --- |
| **Data collection and processing** |  |
| Magnification | 105,000 |
| Voltage (kV) | 300 |
| Electron exposure (e<sup>-</sup>/Å<sup>2</sup>) | 56.15, 58.32 |
| Defocus range (μm) | -1- -2 |
| Pixel size (Å) | 0.85 |
| Symmetry imposed | C1 |
| Initial particle images (no.) | 6,687,338 |
| Final particle images (no.) | 1,006,848 |
| Map resolution (Å) | 3.0 |
| FSC threshold | 0.143 |
| Map resolution range (Å) | 2.5-4.5 |
| **Refinement** |  |
| Initial model used (PDB code) | 7S8N, 7F4H |
| Model resolution (Å) | 3.0/3.3 |
| FSC threshold | 0.143/0.5 |
| Model composition |  |
| Non-hydrogen atoms | 9679 |
| Protein residues | 1232 |
| Ligands | 1 |
| B factors (Å<sup>2</sup>) |  |
| Protein | 94.81 |
| Ligand | 158.10 |
| R.m.s. deviations |  |
| Bond lengths (Å) | 0.003 |
| Bond angles (°) | 0.660 |
| Validation |  |
| MolProbity score | 1.81 |
| Clashscore | 8.11 |
| Poor rotamers (%) | 0.19 |
| Ramachandran plot |  |
| Favored (%) | 94.65 |
| Allowed (%) | 5.35 |
| Disallowed (%) | 0 |

